# Experiments and modelling of pulmonary surfactant disruption by aerosolised compounds

**DOI:** 10.1101/2024.10.16.618442

**Authors:** Hugh Barlow, Sreyoshee Sengupta, Maria Teresa Baltazar, Jorid B. Sørli

## Abstract

Within the deep lung, pulmonary surfactant coats the air-liquid interface at the surface of the alveoli. This complex mixture of amphiphilic molecules and proteins modifies the surface tension and mechanical properties of this interface to assist with breathing. In this study, we examine the effects on pulmonary surfactant function by two industrially used compounds composing surfactants and polymers. Using an experimental method previously developed to imitate the in vivo exposure in the alveoli[1], we quantify the change in the dilational rheology of the pulmonary surfactant due to the introduction of two widely used chemicals; Benzalkonium Chloride (BAC) and Polyhexamethylene Biguanide (PHMB). We observe that these chemicals alter the dilational rheology of the surfactant monolayer. Using a mechanistic theory, we are able to semi-quantitatively model the changes induced by the introduction of these compounds to the pulmonary surfactant.

## Introduction

Human pulmonary surfactant is a complex mixture of phospholipids and proteins that lines the air-liquid interface within our alveoli. Its composition and functionality is a topic of great interest from the perspective of both fundamental colloidal science [2, 3] and human health [4]. It is predominantly (>90%) composed of phospholipids, with much of the remainder made up of specialised surfactant proteins. During expiration, it allows the surface tension to reach near zero and despite the extremely high surface pressure does not undergo collapse. This is due to the interplay between the various surfactant proteins which regulate the migration of the surfactant from the surface to multilayer subsurface structures. Upon inhalation, and the corresponding expansion of the alveoli, pulmonary surfactant respreads much more quickly than would be observed in pure dipalmitoyl phosphatidylcholine (DPPC) monolayers[2]. This rapid respreading overall increases the elasticity of the lung which is critical to maintaining respiratory function. It is noted in an early study “that it is high surface elasticity rather than very low surface tension which is decisive for normal lung function”[5].

Many studies of the pulmonary surfactant have studied these rheological properties and their importance to healthy breathing [3, 6-9]. Researchers have analysed how pulmonary surfactant monolayers can be altered by the interaction with non-biological materials (xenobiotics) such as nanomaterials [10-17], e-cigarette components [18, 19] and polymers [1, 20-22]. The delicate interplay between the rheology of the lung and human respiratory health suggests that the induced change of the monolayer properties could induce an adverse response in persons exposed to these aerosolised compounds. This was notably seen in the case of water-proofing polymer-based sprays[23, 24]. Zasadzinski and co-authors have suggested a link between a change in the dilational modulus and Acute Respiratory Distress Syndrome [7, 25, 26]. They have suggested that such a decrease may lead to a “Laplace instability”, whereby the alveoli fail to inflate and deflate in accordance with normal breathing due to a decrease in the surface elasticity compared to the average surface tension, potentially causing fatal respiratory failure.

It is therefore of tremendous importance to public health that the potential impact of chemicals on lung surfactant function is assessed. Increasingly, regulatory and consumer pressures have lead to the adoption of non-animal methods to assess the risk from novel chemicals and products[27]. We study here the alteration in the interfacial properties of lung surfactant using in vitro experiments and theoretical modelling derived from well established surfactant physics. We use two chemicals: a cationic surfactant benzalkonium chloride (BAC) and a hydrophilic polymer polyhexamethylene biguanide (PHMB) as model compounds. BAC is safely used in various consumer (e.g. cleaning products) and pharmaceutical products (e.g. nasal sprays and ophthalmic drops) due to its broad-spectrum antimicrobial activity [28]. However, in the asthmatic population, nebulisers containing BAC have resulted in cases of bronchospasm and bronchoconstriction [29]. These nebulizers generate small particles (2-5 µm) are designed to deliver pharmaceutical drugs mainly to the distal bronchial region but can also reach the alveolar area[30]. This contrasts with the particle size distribution produced by trigger sprays used in disinfectants, which generate median particles from about 70 µm up to well over 100 µm[31]. These are unlikely to reach the tracheobronchial area [32].

PHMB is used as a preservative in cosmetic products at a concentration up to 0.1% as evaluated by the Scientific Committee on Consumer Safety (SCCS). Based on the severity and irreversibility of the lung effects observed in the acute and subacute inhalation toxicity animal studies at low concentrations (No observed adverse effect level of 0.02 mg/m^3^), and in the absence of sub-chronic inhalation studies, use in sprayable formulations is not advised[33]. PHMB is closely related to other anti-microbial polyhexamethylene based polymers which have a known pulmonary health impact when used in aerosols[34].

## Experimental Method

The experimental setup used in this study is diagrammed in Figure 1. The method used has been outlined in detail in previous work[1, 21, 22, 35-41], so we shall only describe it briefly here. The Constrained Drop Surfactometer (CDS, BioSurface Instruments, United States) was used to measure the surface tension of pulmonary surfactant under dynamic conditions,[17]. The CDS setup was used with a constant flow through exposure chamber as described in [37] to study the effect of aerosolised test chemicals on pulmonary surfactant function.

**Figure 1:**
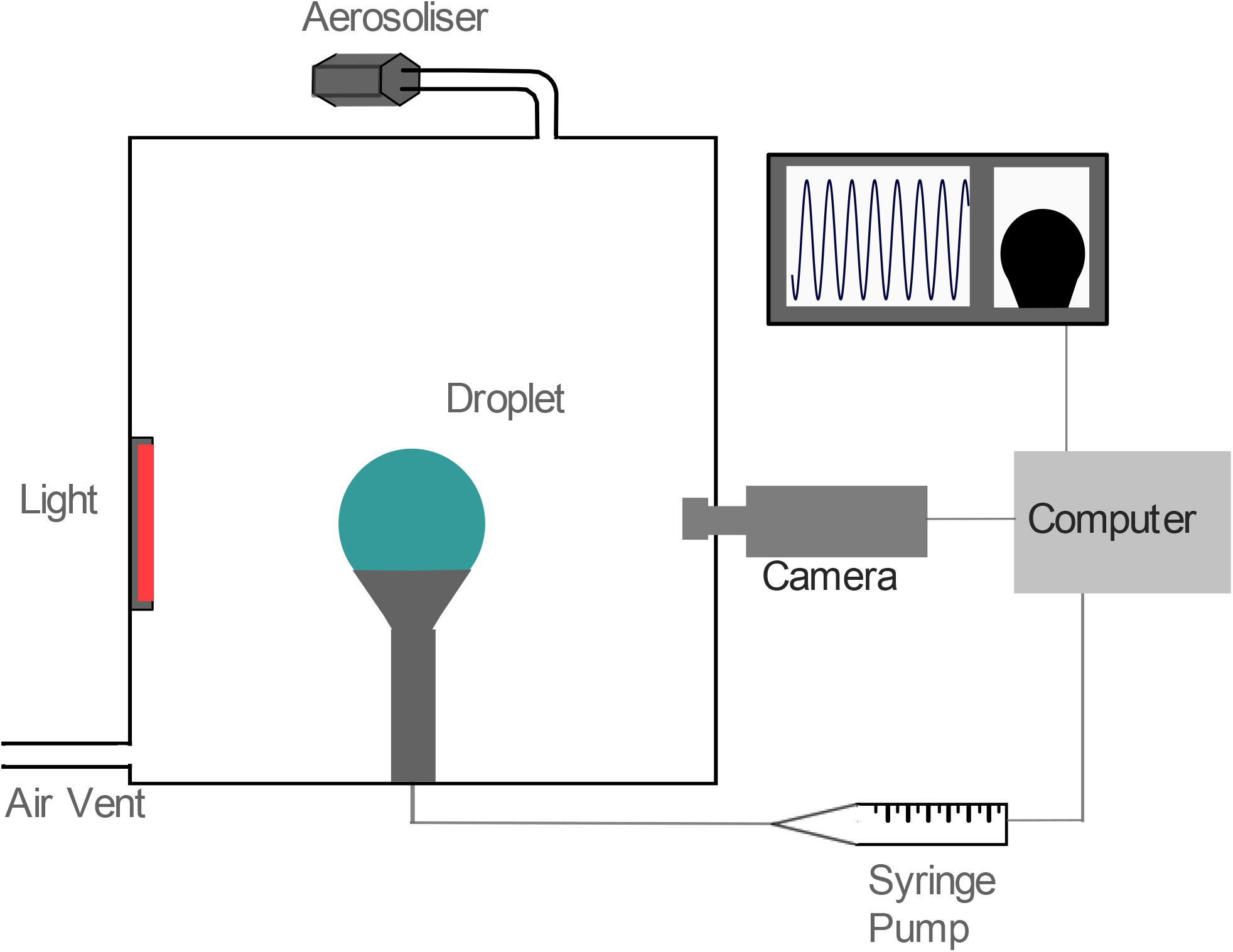
Schematic of the experimental setup with CDS, mechanised pump, camera and computer reading of surface tension.

A drop of commercially purchased pulmonary surfactant (Curosurf™, 10 µL at 2.5 mg/mL) was placed on a hollow, sharp-edged pedestal connected via plastic tubing to a syringe filled with buffer, the syringe was placed in a computer controlled motorised pump. The drop was subjected to dynamic cycling (20 cycles/minute, at 20-30% compression) by introducing and removing liquid from the drop through the base of the pedestal to simulate breathing. The cycling of the drop was stopped to refill the drop with buffer and replace the evaporated liquid when the drop became too small.

Pulmonary surfactant was cycled for 40 seconds to obtain a baseline value followed by exposure to the aerosolised chemicals for 5 minutes. Experiments with a baseline minimum surface tension value >5mN/m or compression >30% were discarded. Drop images were continuously recorded at five frames per second and analysed by axisymmetric drop shape analysis (ADSA) [42] software to calculate surface tension and surface area among other parameters. The temperature in the exposure chamber was monitored using the TinyTag Plus 2 data logger (TGP-4017, Gemini Data Loggers Ltd, United Kingdom). The CDS setup was placed in a heating box and the air used to aerosolise the test chemical was heated. Aerosols of the test chemical were generated by infusing a solution of chemical in solvents (ethanol for BAC and water for PHMB) with an infusion pump (Legato 100, KD Scientific, Holliston, USA) into a Pitt no.1 nebuliser [43] by the flow of warm pressurized air into the nebulizer. Data from the ADSA software was processed to quantify changes in surface tension.

### Rheological Analysis

In this study, we seek to examine how the dilational rheology of pulmonary surfactant changes when exposed to an aerosolised compound during dynamic cycling in constrained droplet surfactometer (CDS). This is done by determining the complex dilational modulus *E* using the equation

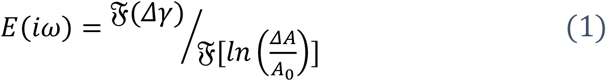

where 𝔉 denotes the Discrete Fourier Transform operator, ln (*ΔA*/*A*_0_) the relative logarithmic surface area, with circular frequency *ω*, and *γ* is the surface tension. Similar processes have been used in the past to study the dilational rheology of surfactant monolayers [44-47], including for pulmonary surfactant dilational rheology[10, 11].

In Figure 2 a) we plot an example of the surface tension *γ* as function of time observed in our experiments. This example corresponds to BAC at an infusion rate of 0.1 ml/min and concentration of 0.5 mg/ml. Prior to the exposure time, when the chemical is infused into the chamber, a steady oscillation is observed, the amplitude of which decreases precipitously upon introduction of the aerosolised xenobiotic. This infusion is conducted continuously until the end of the measurement time at 300 seconds. We note that the new post-exposure amplitude of oscillation does not alter much over time after the initial transition from the pre-to post-exposure regime. However, there is a non-negligible transition time between the pre- and post-post exposure amplitude, and taking measurements over the entire post exposure duration may introduce artifacts to our measurements due to this intermediary phase. Furthermore, during these experiments, the solvent in the droplet fluid is constantly undergoing evaporation which can alter the measured surface tension due to resulting increase in solute concentration in the droplet. Consequently, we perform the measurements to characterise the pre- and post-exposure dilational rheology in sub-periods of the pre and post exposure regimes. These sub-periods are selected where the variation in the surface area and surface tension of the droplet most closely correspond the single frequency sinusoidal oscillation applied through the apparatus, i.e. where the surface-tension and surface area variation with time can be best characterised by a single Fourier mode. To obtain decent averaging these subperiods are required to correspond to not less than 10 oscillations (30 seconds) in duration. These sub-periods are shown in the Figure 2 a) for the pre-exposure and the post-exposure sub-periods as green and pink solid lines respectively. Further experimental explorations using more precise equipment and longer exposure times using methods to limit evaporation may resolve these issues in future.

**Figure 2:**
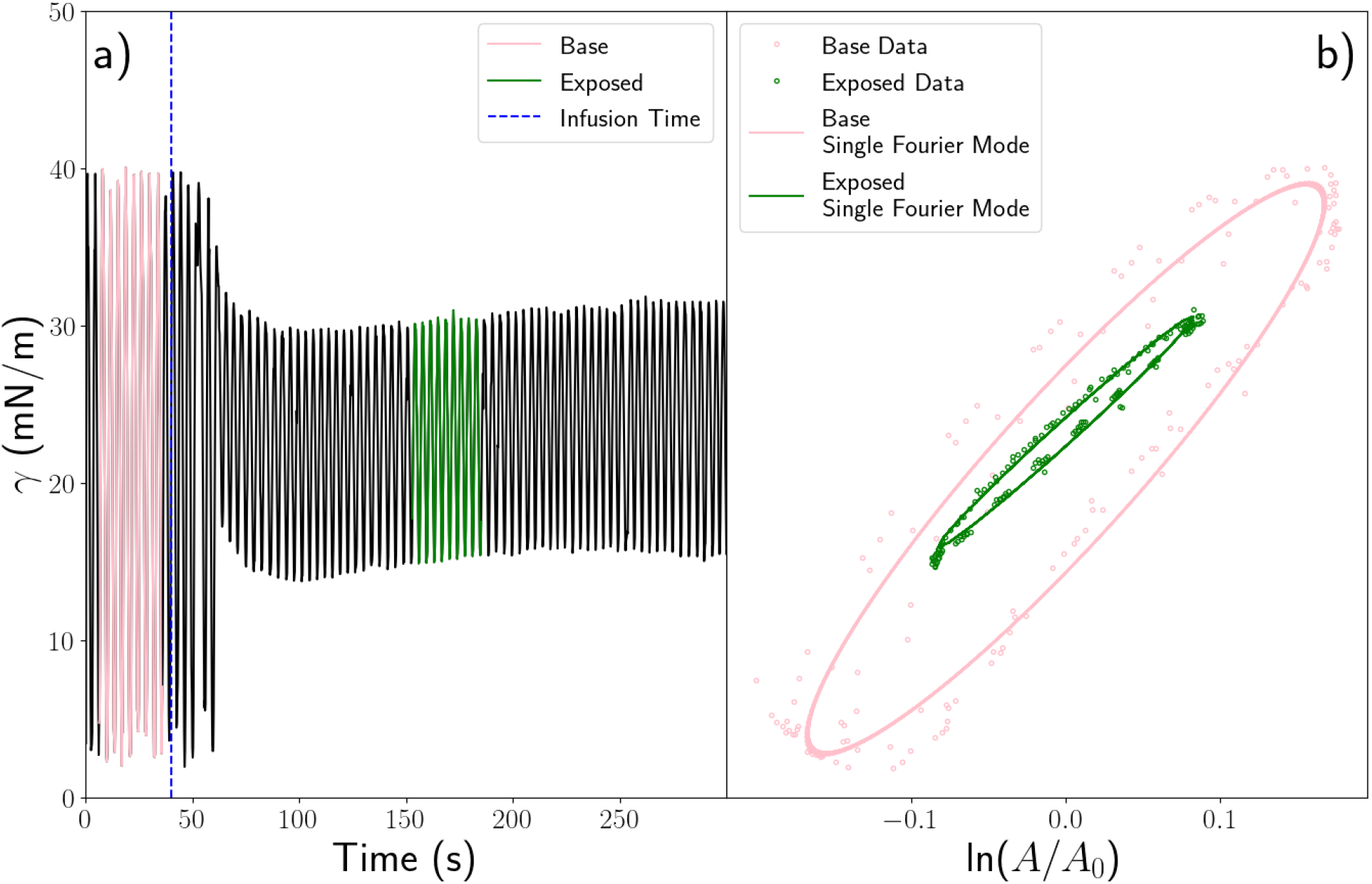
a) Surface Tension as a function of time measured in experiment. Infusion rate of BAC is 0.1 ml/min and concentration of 0.5 mg/ml. Infusion begins at 40s. b) Lissajous curves corresponding to the unexposed and exposed parts of the time course data in a) Data (·) and theoretical Lissajous curve corresponding to the largest Fourier mode (-).

In Figure 2 b), we show the corresponding parametric plots (Lissajous Curves) of these sub periods, where the coloured symbols correspond to the data from the experimental data shown in Figure 2 a).

Overlaid in solid lines are the elliptical Lissajous curves given by a parametric plot of the largest measured Fourier mode of the surface tension against surface area added to the average surface tension. The equation of these curves is given by

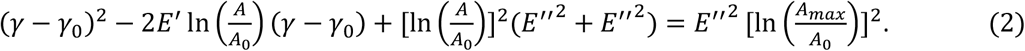

Here *E*^′^ and *E*^′′^ are the storage and loss dilational modulus respectively, *γ*_0_ is the average surface tension, *A*_0_ is the average droplet area and *A*_*max*_ is the largest droplet area during oscillation. The corresponding complex modulus is then *E*^∗^ = *E*^′^ + *i E*^′′^· We see that the elliptical curve approximates very well these curves before and after exposure. We therefore shall use the moduli calculated based on the largest amplitude Fourier mode to characterise the rheology before and after the introduction of the xenobiotic.

We quantify the observed change in the viscoelastic properties of the pulmonary surfactant as a function of the introduced compound by examining the change in the complex modulus *E*^∗^ and the average surface tension 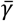.We define the normalised difference in the complex dilational modulus 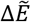 as

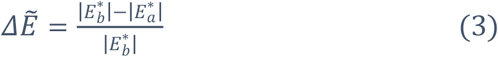

where *E*^∗^_*b*_ and 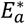 are the complex moduli before and after the exposure of the droplet to the chemical and 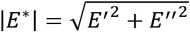 is the magnitude of complex modulus *E*^*^. The corresponding normalised change in the average surface tension 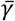 is quantified by the expression,

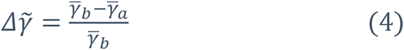

Where 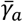 and 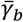 are the average surface tension before and after exposure is initiated. In both cases, we normalise the change to minimise the impact of sample variability in the pulmonary surfactant base behaviour that may be result from inconsistency in experimental material.

## Modelling

We now present a model to understand the effects of the xenobiotic on the native pulmonary surfactant in our experiments based on the physics of surfactant monolayers. The dilational rheology of pure surfactant monolayers is predominantly determined by the kinetics of molecular migration between the bulk fluid and the interface[48]. Due to the interactions between the phospholipids and proteins that compose pulmonary surfactant, the changes in surface pressure observed therein during dynamic compression is more complex[6, 8]. This has been linked to the formation of three-dimensional structures beneath the monolayer which act as reservoirs of surfactant molecules during decompression. [3, 49-53]. These regimes of behaviour are not accounted for by the kinetics of adsorption and desorption[3]. Multiple studies have been performed that developed mathematical models which encompass these different behaviours seen during the dynamic compression of pulmonary surfactant monolayers [54-58].

In this study, we do not seek to comprehensively model comprehensively the changes in surface tension seen during the dilation of pulmonary surfactant monolayers in all regimes, but to investigate the mechanisms related to the disruption of pulmonary surfactant function. We therefore build a minimal model to describe the dynamics of the native pulmonary surfactant based on the dynamics of surfactant adsorption and ignore the behaviour at higher levels of compression. As in Ref.[6], we relate the surface concentration of pulmonary surfactant Γ to the surface pressure П using a Volmer type isotherm[59]

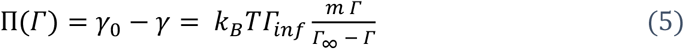

where *k*_*B*_ is Boltzmann’s constant, T is the temperature (assumed 37 °*C*), *γ* is the surface tension, *γ*_0_ is the surface tension of water at zero surfactant concentration and m is an empirical scaling parameter which accounts for the additional elasticity attributable to interactions between surfactant proteins and phospholipids. *Г*_∞_ is the surface concentration at saturation, which is set from the findings of previous studies to 0.02Å^−2^[57, 60, 61]. As in previous studies [54-57], we describe adsorption and desorption by the implied kinetics of the corresponding adsorption isotherm

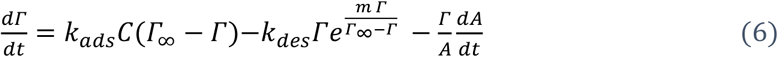

The final term in this equation accounts for the variation in the surface area of the droplet during the oscillating cycles[54, 57]. The parameters *k*_*ads*_*C, k*_*des*_ and *C* correspond to the surfactant adsorption and desorption rate constants and the bulk surfactant concentration respectively. Since *C* does not appear independently, *k*_*ads*_*C* and *k*_*des*_, along with the factor m, are determined by fitting the Equations 4 and 5 to compression cycles of unexposed pulmonary surfactant.

These parameters were obtained by fitting Lissajous curves for 10 different samples of commercially available lung surfactant Curosurf™ at a fixed oscillation rate of 20 *min*^−1^ to emulate human breathing frequency. In Figure 3 a) we plot one of these experimental Lissajous curves and the model curve used in this study and we see good agreement between the model and the experimental data. The fitted parameters used throughout this study are shown in Table 1.

**Figure 3:**
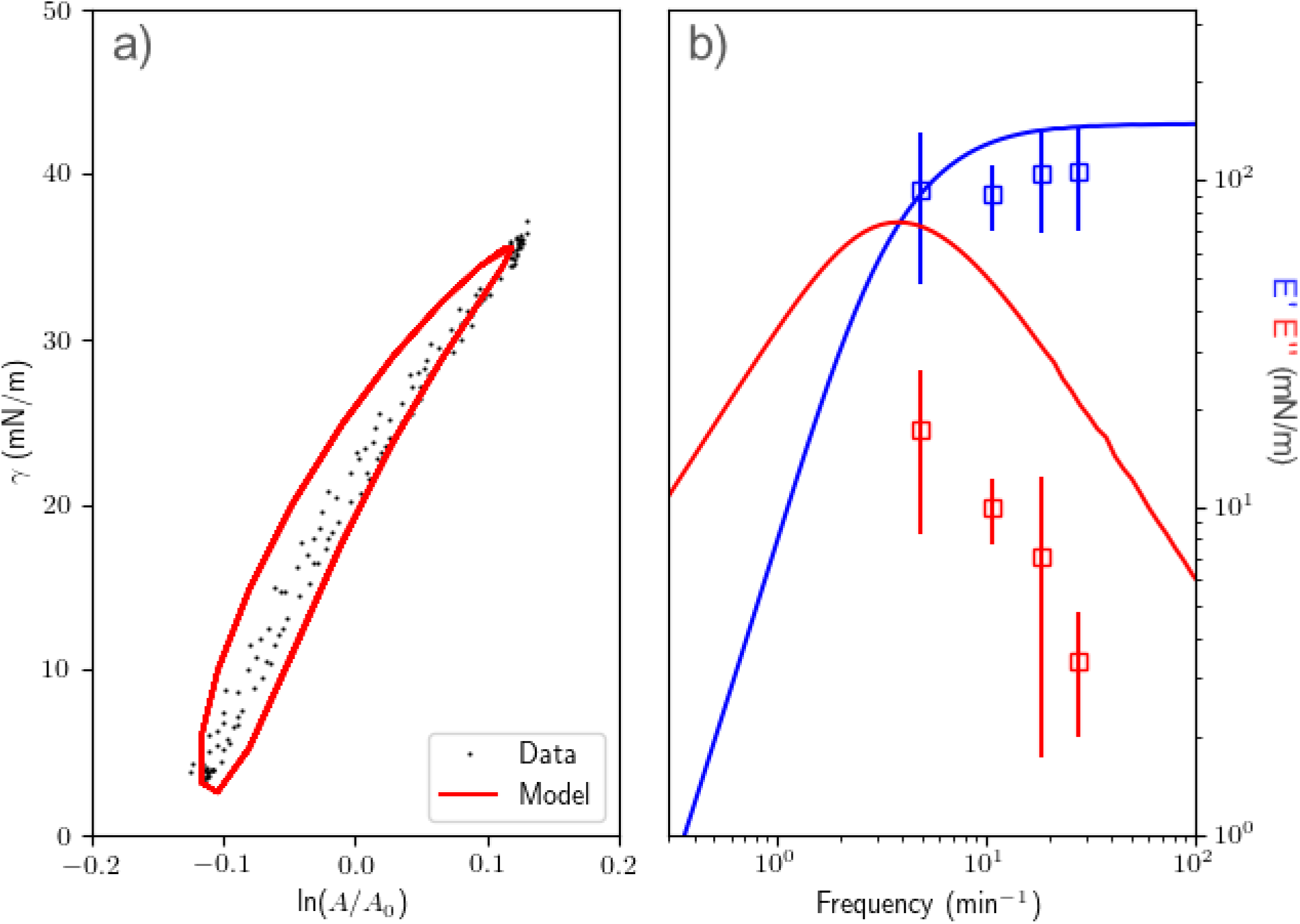
a) Example Lissajous curve of Pulmonary Surfactant at ω = 20 min^−1^, with corresponding model fit. b) Experimentally measured storage modulus E’ and loss modulus E’’ for model pulmonary surfactant based on the largest Fourier mode during oscillatory experiments (□). Model values (-) calculated by applying an amplitude of ln(A/A_0_)_max_ = 0·125· The model values are calculated based on the parameters from best fit parameters for ω = 20 min^−1^. Error bars correspond to one standard deviation of the measured values.

**Tabel 1:** Parameters of best fit for Equation 1 and 2 based on fitting 10 Lissajous curves produced for unexposed lung surfactant.

| Parameter | m | $k_{ads}C$ | $k_{des}$ | $\Gamma_{\infty}$ | $\gamma_0$ |
| --- | --- | --- | --- | --- | --- |
| Units | # | $s^{-1}$ | $s^{-1}$ | $\text{\AA}^{-2}$ | mNm |
| Value | 3.11 | 0.045 | 2.09e-05 | 0.02 | 71.2 |

The measured values for the storage and loss moduli for multiple frequencies are shown in Figure 2b). We see from the plotted curves that our model, although only fitted at a frequency of 20 *min*^−1^ does reasonably well at estimating the storage modulus but overestimates the loss moduli for all frequencies measured. This discrepancy is potentially due to the lack of consideration of the additional mechanisms which govern the behaviour of pulmonary surfactant during periodical oscillation such as subsurface structure formation. However, our model captures the saliant features of the experimental Lissajous curves, therefore we shall use it to model alteration in the pulmonary surfactant oscillatory curve due to the introduction of the xenobiotics studied.

### Xenobiotic Effects

The primary goal of the study is to understand how the introduction of the xenobiotic chemical induces pulmonary surfactant function inhibition. Having demonstrated that we can capture the relevant behaviours of the unexposed pulmonary surfactant using the model above, we now extend it to account for the effects of the xenobiotic by following a methodology similar to that developed to model for the rheology of mixed protein-surfactant monolayers [62, 63].

Firstly, we observe in experiments that when the aerosolised chemical is infused into the chamber, after a short transient period, the amplitude of the oscillations of surface tension settles into a new regular amplitude (see Figure 2 a)). This implies that the xenobiotic does not accumulate on the surface of the droplet but that it is eventually absorbed into the bulk. Therefore, we surmise that the amount of the xenobiotic present in the monolayer is dependent on the rate at which it is deposited as opposed to the total cumulative amount. We set the *dose rate=concentration* ×*infusion rate*. This assertion is corroborated by animal experiments, where it was noted that mice exposed to a low concentration of aerosolised chemicals did not experience deleterious effects [37, 64]. This assertion is tested further for a range of chemicals in Ref. [65] and is found to be generic. In our model, the xenobiotic is introduced to the droplet surface at a fixed rate 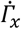. The bulk concentration of the xenobiotic is assumed to remain sufficiently small that no xenobiotic is reabsorbed to the surface from the bulk, i.e the droplet bulk is considered as a sink. The desorption kinetics are assumed to be described by the same kinetics as the pulmonary surfactant molecules, albeit with a different desorption rate constant *k*_*x,des*_.

In addition to the short-range interactions described by the Volmer isotherm (Eqs. 5 and 6), we consider the influence of longer-range interactions between molecules. We must therefore include the free energy of interaction between the lung surfactant and the xenobiotic which are up to third-order *F*_*int*_ = *β*_1,*x,ps*_ *Г*_*x*_*Г*/*Г*_∞_ + *β*_2,*x,ps*_*Г Г*_*x*_^2^/*Г*_∞_^2^, where *β*_1,*x,ls*_ and *β*_2,*x,ls*_ are the xenobiotic-pulmonary surfactant and xenobiotic-pulmonary surfactant interaction parameters corresponding to second and third order interactions respectively. In Ref.[63], it was assumed that the second-order mean field interaction energy is zero (*β*_1,*x,ls*_ = 0) due to the formation of condensed phases of protein in the combined protein surfactant monolayer. If *β*_1,*x,ps*_ ≠ 0, the kinetics will include the term *β*_1,*x,ls*_(*Г*_*x*_ + *Г*)/*Г*_∞_ in the exponent which is non-zero at the limit of *Г*_*x*_ → 0, for non-zero *Г*, leading to non-physical behaviour. We therefore shall also set *β*_1,*x,ls*_ = 0 for our model. This leads to the exponential term in the kinetics of *β*_2,*x,ls*_*Г*_*x*_(*Г*_*x*_ + *Г*)/*Г*_∞_^2^ which vanishes in absence of the xenobiotic, reflecting the observations in the performed experiments. For clarity, hereafter we shall omit the subscript on this interaction parameter and the *β* shall be used to refer to *β*_2,*x,ps*_ throughout.

It is experimentally infeasible to determine the exact value of 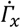 (the average amount deposited on the droplet per unit time). Therefore, it is necessary to introduce a scaling parameter α, such that the actual infusion rate is proportional to the dose rate, 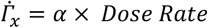. As a result, the change in the saturation concentration *Г*_∞_ is obfuscated, since we cannot discern the exact number of molecules deposited. Therefore, we assume that the value of *Г*_∞_ remains unchanged. A future study which includes a precise measurement of the deposited mass would address this.

Combining these assumptions, we obtain the following equations that describe the surface pressure and kinetics of the combined xenobiotic and native pulmonary surfactant system during exposure;

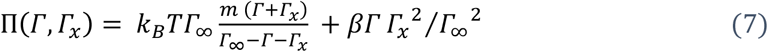

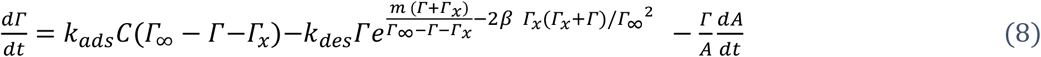

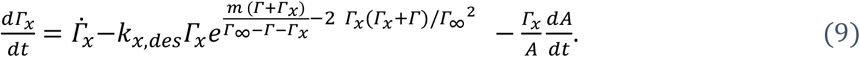

Using these three equations we aim to model the changes in average surface tension and rheology seen in our experiments. For both chemicals considered in this work, we will fit the three parameters *α,k*_*x,des*_ and *β*·

## Results and Discussion

For both PHMB and BAC, experiments were performed at multiple concentrations and infusions rates. For each combination, multiple repeats were performed. The average of the measured 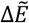 and 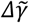 for each combination of dose and infusion rate are plotted in Figure 4 with error bars corresponding the one standard deviation. For both BAC and PHMB we observe a similar trend in 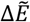,an initially gradual increase with increasing dose-rate, which steeply rises at some critical value.

**Figure 4:**
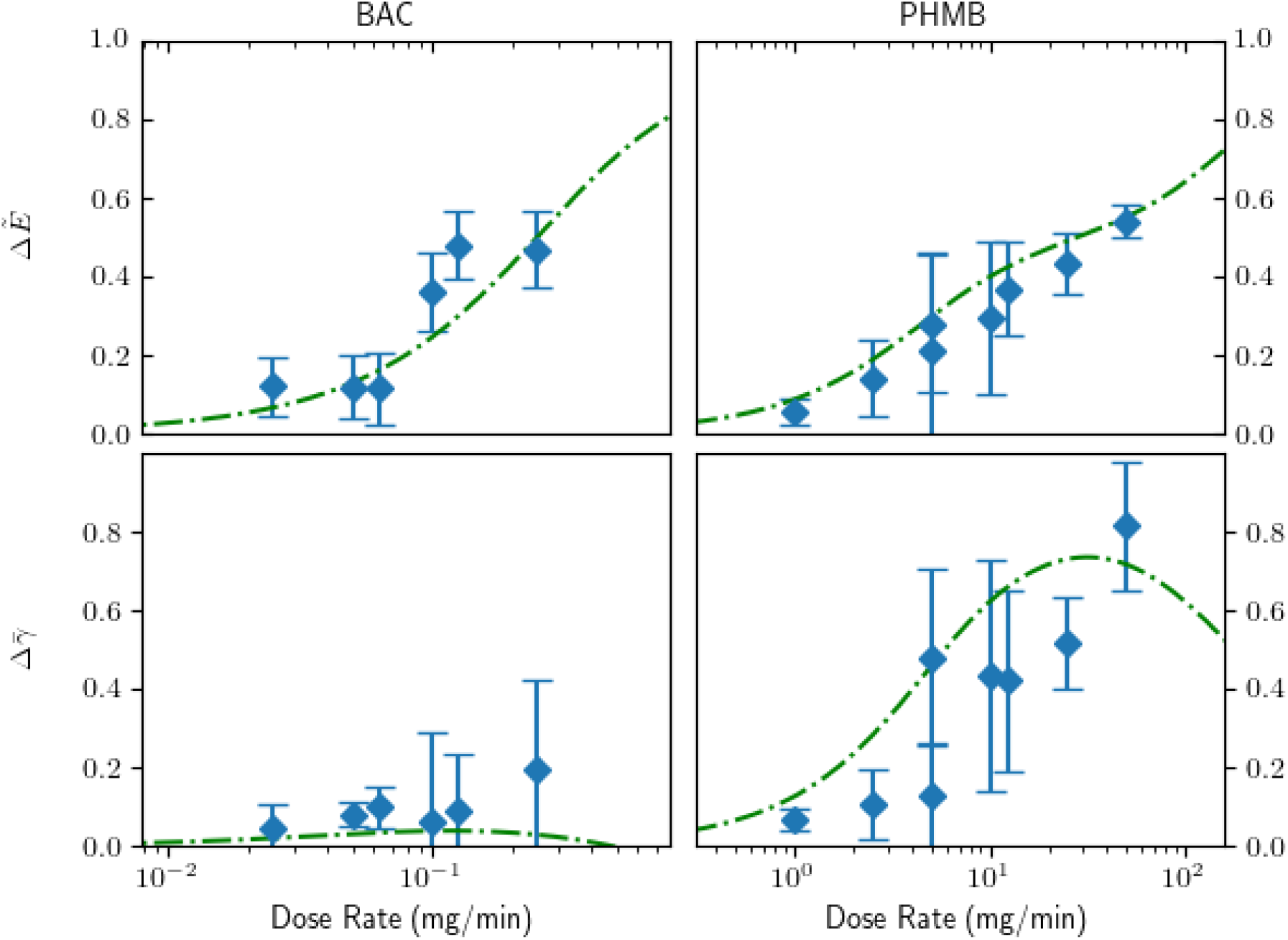
Normalised change in complex dilational modulus and average surface tension. Experimental data shown as symbols and fitted model as lines.

However, a distinct difference is seen with the change in the average surface tension, which increases for PHMB but shows almost no change for BAC. This may be explained by the BAC being an amphiphilic surfactant, whereas PHMB being a hydrophilic polymer will not demonstrate surfactant properties at the surface concentrations in this study.

We then fit Equations 6-8 to the calculated the values of 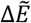 and 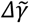.The parameters of best fit are determined using the SciPy optimize library to minimize the difference between these theoretically calculated values and the corresponding experimentally measured results. These best fits are shown as dashed lines in Figure 2 and we see that for both PHMB and BAC the model fits the data reasonably well and the parameters of best fit are shown in Table 2.

**Table 2:**
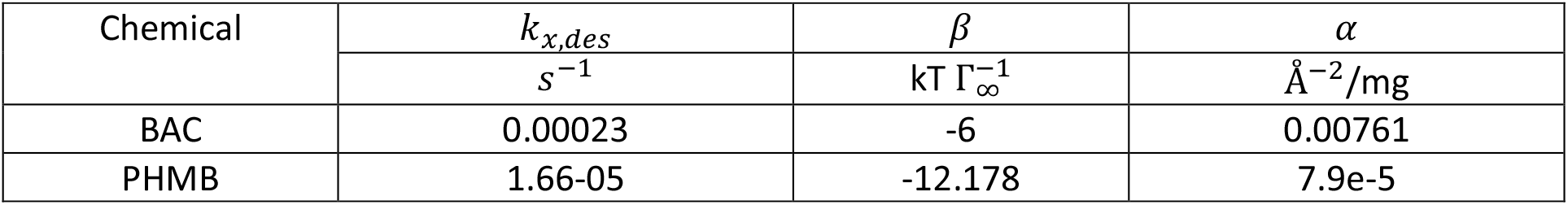
Parameters of best fit for Equation 6-8 for tested compounds.

| Chemical | $k_{x,des}$ | $\beta$ | $\alpha$ |
| --- | --- | --- | --- |
| | $s^{-1}$ | $kT \Gamma_{\infty}^{-1}$ | $\text{\AA}^{-2}/\text{mg}$ |
| BAC | 0.00023 | -6 | 0.00761 |
| PHMB | 1.66-05 | -12.178 | 7.9e-5 |

although the model captures both 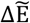 and 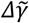 for both models, it predicts that at sufficiently high infusion rates, 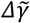 will drop due to the accumulation of material at the interface. Further experiments at higher concentrations and infusion rates could assess this prediction. To look more closely at the behaviour of the model versus the exposed pulmonary surfactant we plot the experimental and theoretical Lissajous curves for PHMB and BAC for multiple dose rates in Figure 5. We see that in both cases there is good-quantitative agreement between the theoretical and experimental curves. However, we do observe that the theoretical curve becomes more circular as the dose rate increases indicating that the viscous modulus *E*^′′^ is increasing relative to the storage modulus in the model, but not in experiments. We note that the unexposed model also overestimates the viscous modulus, so this discrepancy is not unexpected. Despite this, we see reasonably good agreement between the experimental and theoretical curves, suggesting that we have captured the important facets of the induced disruption.

**Figure 5:**
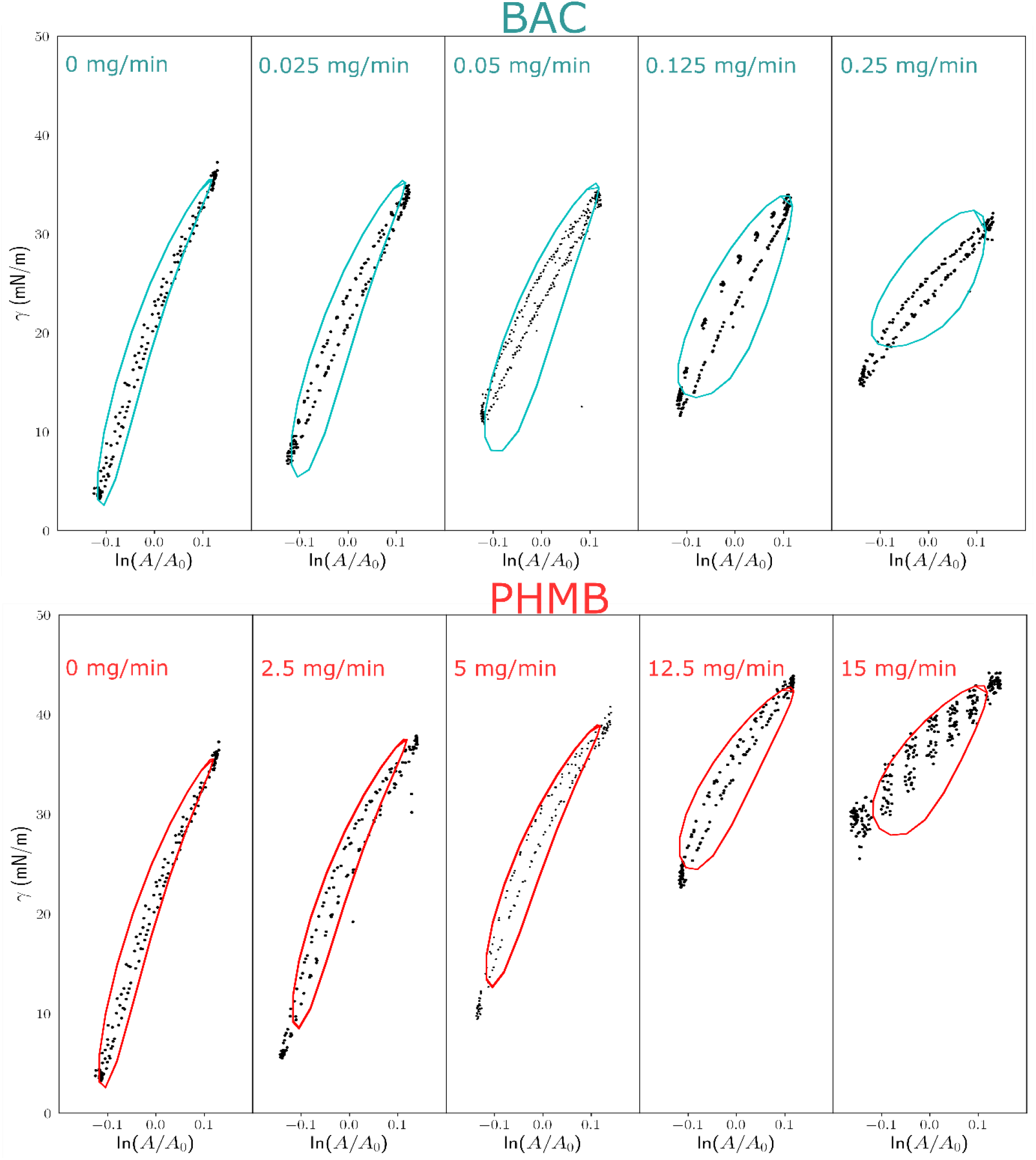
Lissajous curves at different dose rates for BAC (upper) and PHMB (lower) for increasing dose rates. Numbers in each plot correspond to the dose rate for the data depicted.

We now come to consider why some chemicals may induce disruption at a given dose rate and others not. The change in the dilational rheology of the surfactant monolayers is related to the altered desorption kinetics of the monolayer. Both the xenobiotic desorption rate *k*_*x,des*_ and the xenobiotic long range interaction parameter *β*. We may postulate that if *k*_*x,des*_ is sufficiently high (i.e. for an extremely hydrophilic small molecule), the xenobiotic will be adsorbed sufficiently quickly into the bulk. Thereby, only at very high dose rates will enough of the xenobiotic remain on the surface to cause a substantial change in the monolayer composition. This may explain why it has been observed that hydrophobic polymers have been known to induce respiratory distress via pulmonary surfactant disruption when inhaled [21, 22, 64, 66, 67].

We now come to consider why some chemicals may induce disruption at a given dose rate and others not. The change in the dilational rheology of the surfactant monolayers is related to the altered desorption kinetics of the monolayer. Both the xenobiotic desorption rate *k*_*x,des*_ and the xenobiotic long range interaction parameter *β*. We may postulate that if *k*_*x,des*_ is sufficiently high (i.e. for an extremely hydrophilic small molecule), the xenobiotic will be adsorbed sufficiently quickly into the bulk. Thereby, only at very high dose rates will enough of the xenobiotic remain on the surface to cause a substantial change in the monolayer composition. This may explain why it has been observed that hydrophobic polymers have been known to induce respiratory distress via pulmonary surfactant disruption when inhaled [21, 22, 64, 66, 67].

The interaction parameter *β* will also cause a change in the desorption rate of both the xenobiotic and the pulmonary surfactant, further changing the dilational rheology, which again may be the case for a hydrophic material which may interact with the phospholipid tails and hydrophobic surfactant proteins SP-A and SP-B. How this interaction manifests an increased desorption rate remains unknown. One possible method by which disruption occurs is through the formation of micelles of pulmonary surfactant molecules surrounding the xenobiotic which then migrate into the bulk. Discerning this may be done through microscopy or further experiments examining the different regimes of pulmonary surfactant behaviour and the impact of the xenobiotics thereupon.

## Conclusions

Our study has examined multiple aspects of the inhibition of pulmonary surfactant by aerosolised compounds. We have incorporated in vitro experiments, rheometric analysis and theory to how such compounds may induce changes to the rheological properties of pulmonary surfactant monolayers, potentially leading to disruption of normal breathing. In our study, we have observed a dose-rate dependence on the alteration of the pulmonary surfactant properties that corroborates the findings of previous studied where it was noted in Ref. [64] “… that the dose rate rather than the total inhaled dose of substance is critical for the toxic effect”. It was also noted in that study that these results corroborated those seen in human cases where exposure led to hospitalisation.

We have also demonstrated that the salient features of this disruption can be modelled with reasonable accuracy using a model based on surfactant thermodynamics. Our model was able to capture not only the change in the dilational rheology of the lung surfactant, but also the observed change in the average surface tension with reasonable accuracy. However, this modelling does not elucidate what features of the compounds in question drive how they modify the monolayer physics. Further studies, examining chemicals with a variety of physicochemical properties and their impact on lung surfactant functions could determine what features of a compound drive this effect.

We conclude that this work, as well as previous publications [10, 11, 21, 68], has elucidated that respiratory disruption due to certain aerosolised compounds is a physical mechanism as opposed to a biological interaction. In the context of a safety assessment of inhaled materials, this work provides a way to understand the mechanisms of interaction between chemicals and lung surfactant, and the relationship between dose-rate and inhibition of pulmonary surfactant function using computational and in vitro approaches, therefore contributing to the replacement of animal testing.

## Acknowledgements

We are appreciative of the input and help of Iris Muller, Sophie Cable, Joe Reynolds and Anthony Bowden, for fruitful discussions which aided in this work.

## CRediT authorship contribution statement

**Hugh Barlow:** Writing – original draft, Methodology, Formal analysis, Data curation, Visualisation, Software, Investigation. **Sreyoshee Sengupta**: Writing - Original Draft, Investigation, Validation. **Maria Teresa Baltazar:** Supervision, Funding acquisition, Project administration, Conceptualisation, Writing - Review & Editing. **Jorid B. Sørli:** Supervision, Project administration, Conceptualisation, Writing - Review & Editing.

